# bayroot: Bayesian sampling of HIV-1 integration dates by root-to-tip regression

**DOI:** 10.1101/2022.09.20.508733

**Authors:** Roux-Cil Ferreira, Emmanuel Wong, Art F. Y. Poon

## Abstract

The composition of the latent HIV-1 reservoir is shaped by when proviruses integrated into host genomes. These integration dates can be estimated by phylogenetic methods like root-to-top (RTT) regression. However, RTT does not accommodate variation in the number of substitutions over time, uncertainty in estimating the molecular clock or the position of the root in the tree. To address these limitations, we implemented a Bayesian extension of RTT as an R package (*bayroot*), which enables the user to incorporate prior information about the time of infection and start of antiretroviral therapy. Taking an unrooted maximum likelihood tree as input, we use a Metropolis-Hastings algorithm to sample three parameters (the molecular clock, the location of the root, and the time associated with the root) from the posterior distribution. Next, we apply rejection sampling to this posterior sample of model parameters to simulate integration dates for HIV proviral sequences. To validate this method, we use the R package *treeswithintrees* to simulate time-scaled trees relating samples of actively- and latently-infected T cells from a single host. We find that *bayroot* yields significantly more accurate estimates of integration dates than conventional RTT under a range of model settings.

## 1. Introduction

Root-to-tip (RTT) regression is a simple method to locate the earliest point in time in a phylogenetic tree (*i.e*., rooting the tree; Huelsenbeck et al., 2002), to measure the rate of evolution (Drummond et al., 2003), or to reconstruct the divergence times of common ancestors. This method assumes the existence of a strict molecular clock, *i.e*., the rate at which mutations accumulate is roughly constant over time (Bromham and Penny, 2003). Accordingly, the number of mutations should increase linearly over time. Hence, this method is a linear regression of the evolutionary divergence of sequences from their common ancestor against the times when those sequences were observed. The primary input of RTT regression is an unrooted phylogenetic tree with branch lengths measured in units of evolutionary time (*i.e*., the expected number of substitutions per site; Tajima and Nei, 1984), which is the standard output of maximum likelihood methods for reconstructing phylogenies. The tips of the tree representing observed sequences are labelled with sampling times. Thus, RTT becomes an optimization over three parameters: the location of the root in the tree, the time associated with the root (*x*-intercept), and the molecular clock (slope of regression).

RTT has a broad range of applications. Since many viruses have a very rapid rate of evolution, RTT can be applied to sequences collected over a number of months or years. For instance, RTT has recently been used to estimate the origin date and clock rate of SARS-CoV-2 within the first few months of the pandemic (Duchene et al., 2020). We are particularly interested in the use of RTT to estimate the integration dates of HIV-1 proviruses within hosts (Jones et al., 2018). HIV-1 converts its RNA genome into double-stranded DNA that becomes integrated into the host genome as part of the virus replication cycle. In some cases, this integrated provirus becomes reversibly dormant in a transcriptionally-inactive host cell (Siliciano and Siliciano, 2004). This long-lived reservoir of latently-infected cells is the primary obstacle to an effective cure for HIV-1. Consequently, characterizing the composition and dynamics of the latent reservoir has significant implications for HIV-1 cure research (*e.g*., Gondim et al., 2021).

For instance, we can estimate the molecular clock (the slope of the regression) from longitudinal samples of plasma HIV-1 RNA sequences before the start of antiretroviral therapy (ART). If we reconstruct a tree relating both these RNA sequences and proviral sequences from the latent reservoir, we can then use our clock estimate to extrapolate integration dates for the latter (Jones et al., 2018). This relies on the assumption that the integrated HIV-1 genome ceases to accumulate mutations upon integrating into the host genome. Due to its simplicity, RTT has a number of significant limitations. It implicitly assumes that the input tree is known without error. In addition, RTT methods generally yield a single ‘point estimate’ of model parameters by minimizing some cost function (Drummond et al., 2003; To et al., 2016). Mapping proviral sequences to the regression line yields one and only one estimate of the integration date. However, variation in the number of mutations after a given amount of time is expected, even under a strict molecular clock (Langley and Fitch, 1974). A proviral sequence may, by chance, carry more mutations than expected given its actual date of integration. This can cause RTT to project a sequence’s integration date estimate into the future, past its time of sampling or even past the start of ART, when the infection of new cells should be completely suppressed.

Here we describe a Bayesian extension of the RTT method to estimate HIV-1 integration dates. Adopting a Bayesian approach provides a means of quantifying our uncertainty in estimating integration dates, as well as incorporating prior information about the time of infection and the start of ART. We detail our implementation of this method as an R package called *bayroot*, and use a simulation model of within-host population dynamics to validate *bayroot* in comparison to conventional RTT.

## 2. Methods

### Regression model

We start with an unrooted tree *T* relating *n* observed sequences. A strict molecular clock assumes that mutations accumulate at a constant rate *μ* over time, such that the number of mutations per unit time follows a Poisson distribution. Let *Y*_*i*_ be the number of mutations in the *i*^*th*^ observed sequence, which is determined by the location of the root in *T*. Since *Y*_*i*_ is an integer-valued outcome, we must rescale the input tree *T* by multiplying its branch lengths by the sequence length, such that lengths are in units of the expected number of substitutions per genome. Let *t*_0_ be the origin time associated with the root. Let Δ*t*_*i*_ be the time that has elapsed between the *i*^*th*^ sample and the root. The log-likelihood for a set of RNA sequences {*Y*_*i*_, Δ*t*_*i*_} is:

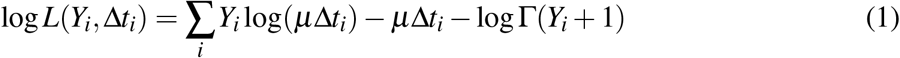

where Γ(*x*) is the gamma function. Equation (1) is sometimes referred to as the Langley-Fitch model (Langley and Fitch, 1974).

We assume a uniform prior distribution for possible locations of the root over the entire length of the tree. We also assume a uniform prior distribution for *t*_0_. If a seroconversion window, *i.e*., the time interval between the last HIV seronegative visit and the first seropositive visit, is available for the host individual, these visit dates can be used to set lower and upper bounds for the uniform prior. Finally, we assume a lognormal prior distribution on the clock rate *μ*, which can be informed by previous measurements of HIV-1 substitution rates within hosts (*e.g*., Alizon and Fraser, 2013).

With these prior distributions and the model likelihood, we implemented a Metropolis-Hastings sampling algorithm in R. A proposal function shifts the root along a branch by some distance *δ*, selecting a branch at random if it encounters an internal node, *i.e*., split, as it traverses the length of the tree. If, however, a terminal node is encountered before the root has been shifted by distance *δ*, then the remaining distance is traveled by reflecting back from this terminus. This results in a symmetric proposal distribution. We also used a uniform proposal *μ*′ *∼* Unif(*μ − δ, μ* + *δ*) for the clock rate, and a truncated normal proposal 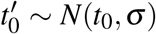 for the origin time. The sampling algorithm returns an S3 object storing a data frame of sampled parameter values and a character vector of sampled trees serialized into Newick strings.

### Sampling integration dates

Given a posterior sample of parameters *Y, μ* and *t*_0_, we need to propagate this information to the distribution of integration times associated with DNA sequences sampled post-ART initiation. Using Bayes’ rule, the probability of integration time *t* _*j*_ for the *j*^th^ HIV-1 DNA sequence given divergence *Y*_*j*_ is:

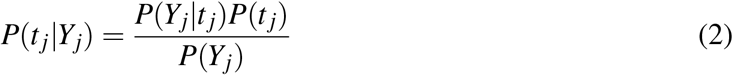

where we index by *j* instead of *i* to emphasize a shift from RNA to DNA sequences. We assume a uniform prior for integration times, *P*(*t*_*j*_) = (*T −t*_0_)^*−*1^, where *t*_0_ is the origin date and *T* is the time of ART initiation. Substituting equation 1 and setting *s* = *t −t*_0_, we solve the integral *P*(*Y*_*j*_) in the denominator as:

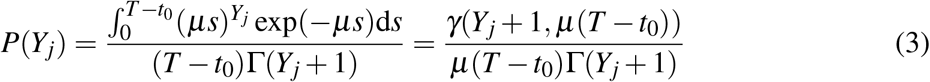

where *γ*(*a, x*) is the lower incomplete gamma function, 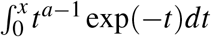. Finally, substituting equations (1) and (3) into (2) and letting Λ = *μ*(*T −t*_0_), we can write:

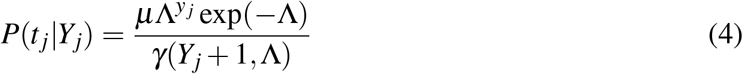

To generate a sample of integration dates, we use a simple rejection sampling method. For a given posterior sample of *Y*_*j*_, *μ* and *t*_0_, we use Brent optimization to locate the maximum of Equation (4), initialized at the midpoint *t* = *t*_0_ + (*T −t*_0_)*/*2. This maximum was used as an upper bound for rejection sampling for values of *t ∼* Unif(*t*_0_, *T*).

The Bayesian regression and integration date sampling methods described above were implemented in R as a package called *bayroot*. All source code is publicly available under the MIT license at https://github.com/PoonLab/bayroot.

### Simulating data

To validate the above method, we used the R package *twt* (‘trees within trees’, https://github.com/PoonLab/twt) to simulate cell population dynamics forward in time, and then to simulate trees by sampling lineages backwards in time to their common ancestors. This package uses the exact stochastic simulation of discrete events (Gillespie, 1977). In brief, it calculates the total rate of all events (Λ), draws an exponentially distributed waiting time to the first event *τ ∼* exp(*−*Λ), and then draws a uniform random number to determine which event occurs. We implemented a compartmental model of cell population dynamics (Figure 1) that can be represented by the following set of differential equations:

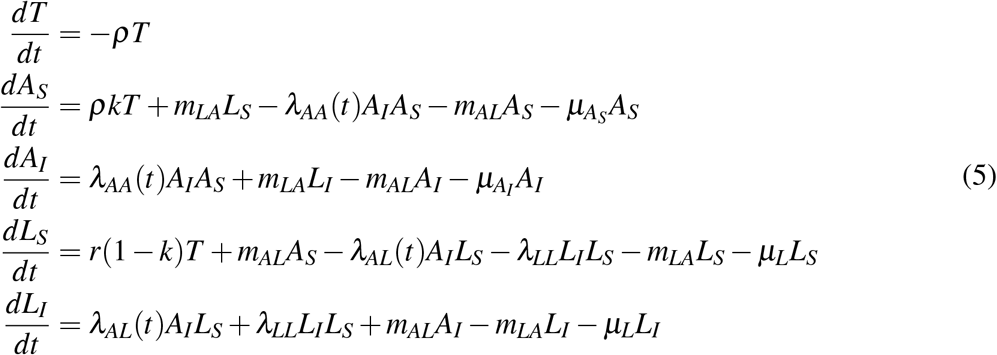

**Figure 1:**
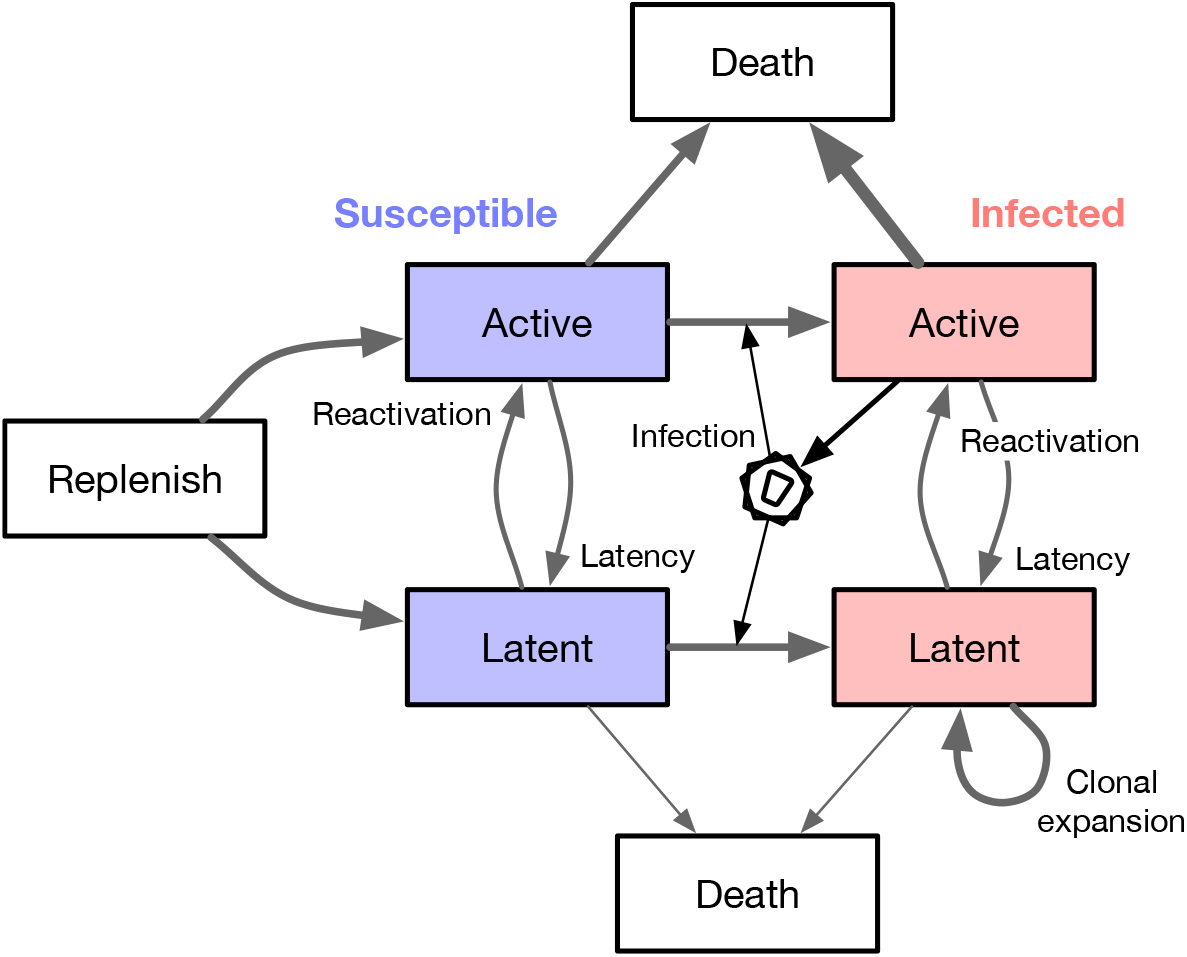
A schematic diagram of the compartmental model used to simulate cell population dynamics. Each box represents a well-mixed population of cells sharing the same rate parameters. We assume that only actively-infected cells release virus particles that go on to infect other, susceptible cells.

This model is a simplified version of the system described by Rong and Perelson (2009). Most notably, our version does not model changes in the viral load. *T* represents a finite population of naive CD4+ T cells from which the populations of active (*A*) and resting (latent, *L*) cells are replenished at rates *kρ* and (1 *− k*)*ρ*, respectively, for 0 ≤ *k* ≤ 1. The *S* and *I* subscripts denote susceptible and infected subpopulations of active and latent cells. A branching event (*λ*_*xy*_) requires a source cell to induce a target cell to undergo a change of state (switch compartments from *x* to *y*). For example, *λ*_*AA*_ represents the infection rate of a susceptible active T cell by a virus released from an actively infected cell. We assume that virus replication is completely blocked by the initiation of ART at time *t*^*\**^, such that *λ*_*A•*_(*t* ≥ *t*^*\**^) = 0. A transition event occurs when a cell spontaneously migrates from compartments *x* to *y* at rate *m*_*xy*_. For example, *m*_*LA*_ represents the reactivation rate of a latent cell. Lastly, we assume constant cell death rates *μ*_*x*_ for each compartment *x*.

The simulation is initialized at time zero with user-specified population sizes of susceptible cells in each compartment, and a single actively infected cell, *A*_*I*_(0) = 1. We simulated the integer-valued population size trajectories *{T, A*_*S*_, *A*_*I*_, *L*_*S*_, *L*_*I*_*}*(*t*) forward in time until a stopping time of *t* = 20 simulation time units. We generated 50 replicate sets of trajectories under two different scenarios by exact stochastic simulation. The rate parameters were set to the following values: *r* = 0.02, *k* = 0.5, *λ*_*AA*_(*t < t*^*\**^) = 0.002, *λ*_*AL*_(*t < t*^*\**^) = 10^*−*4^, *m*_*AL*_ = *m*_*LA*_ = 0.001, 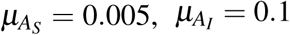, and *μ*_*L*_ = 0.001. ART was initiated at *t*^*\**^ = 10 time units post-infection in scenario 1, and at *t*^*\**^ = 15 in scenario 2. For each iteration of the simulation, we calculated the rates for every type of event, adjusted by the respective compartment size at the current time *t*. For example, the rate of transmissions from *A*_*I*_ to *A*_*S*_ was set to *λ*_*A*_*A*(*t*)*A*_*I*_(*t*)*A*_*S*_(*t*). We drew an exponential waiting time given the total rate of all event types:

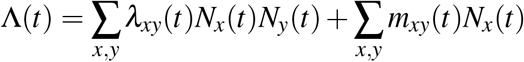

and then determined which event type occurred with probability *λ*_*xy*_(*t*)*N*_*x*_(*t*)*N*_*y*_(*t*)*/*Λ(*t*) or *m*_*xy*_(*t*) *N*_*x*_(*t*)*/*Λ(*t*). Next, we incremented or decremented the respective population sizes for compartments affected by the event type. The time, type and compartments of this event is recorded in a log that is later used to simulate trees. An example set of population size trajectories simulated using this algorithm under scenario 1 is illustrated in Figure 2.

**Figure 2:**
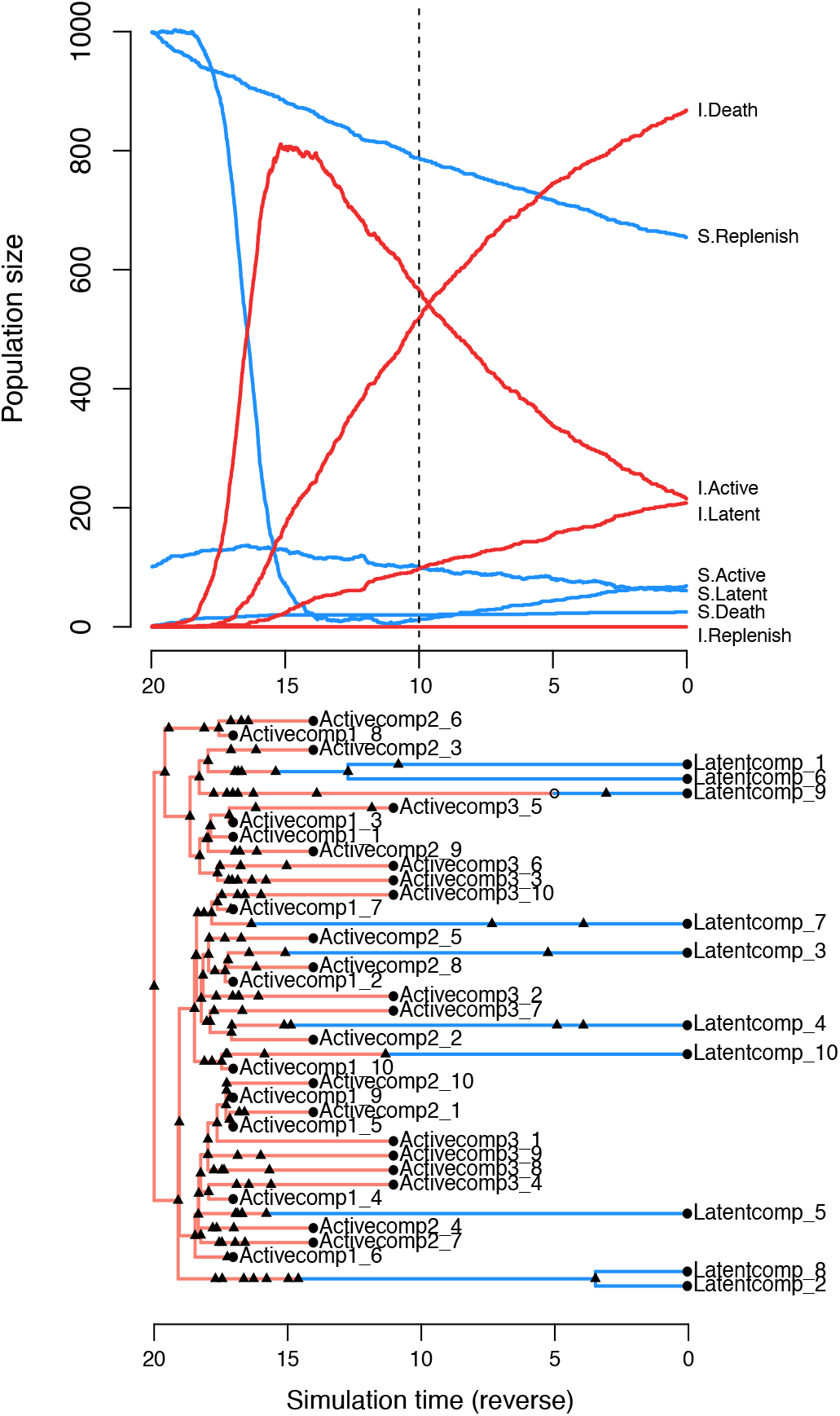
Examples of *twt* simulation outputs for a model of cell dynamics in the latent reservoir. (top) Population dynamics simulated forward in time. Each line represents the population size of a different compartment. S = susceptible, I = infected. The dashed vertical line indicates the time of ART initiation. (bottom) A tree simulated in reverse-time, relating 10 cells sampled from the latently-infected compartment at *τ* = 0, and 30 from the actively-infected compartment at *τ* = 11, 14, 17 (scenario 1), where *τ* = 20 − *t*. Triangles represent transmission events, open circles represent transitions, and closed circles represent sampling times. Branches representing cell lineages in a latent state (blue) are collapsed prior to simulating virus evolution.

To generate a tree relating the sampled lineages in *twt*, we applied another exact stochastic simulation algorithm in reverse time. For the 50 replicate sets of trajectories generated under scenario 1, we sampled 10 HIV-1 RNA lineages at times *t* = 3, 6 and 9 post-infection. For trajectories generated under scenario 2, we sampled 10 HIV-1 RNA lineages at *t* = 11, 13 and 15 post-infection. In both scenarios, we sampled 10 latently-infected cells at *t* = 20 post-infection, for a total of 40 sampled lineages per replicate tree. These lineage sampling times defined the initial conditions for the reverse-time simulation of trees. Next, the algorithm samples events from the log generated in the forward-time simulation to build up a tree relating the sampled lineages. The stopping condition of the tree sampling algorithm is that the sampled lineages converge to a single common ancestor, which becomes the root.

We modified *twt* to output a Newick serialization of this ‘transmission tree’ among cells, labelling tips with sampling times. This tree included internal nodes with only one descendant branch, representing lineage state transitions, or transmissions to/from an unsampled lineage. Internal nodes were labelled with strings encoding the event type, node states (compartments), and unique identifiers for the individual cells involved. These annotations enabled us to ‘colour’ the branches of the tree by lineage state. The true integration dates for sampled latently-infected cells were recorded to a separate file. An example of a tree generated by this process is shown in Figure 2.

To simulate molecular evolution, we collapsed all branches corresponding to latently-infected cells, and used the resulting tree as input for INDELible (version 1.03; Fletcher and Yang, 2009). We assigned an HIV-1 *env* sequence at the root (Genbank accession number AY772699). This sequence is one of the HIV-1 subtype C references curated by the Los Alamos National Laboratory HIV Sequence Database (http://www.hiv.lanl.gov). We configured INDELible to use the Tamura-Nei (TrN) nucleotide substitution model with transition rates *κ*_1_ = 4 and *κ*_2_ = 8, and stationary base frequencies *f*_*A*_ = 0.4 and *f*_*C*_ = *f*_*G*_ = *f*_*T*_ = 0.2. In addition, we rescaled the tree such that the expected number of substitutions per nucleotide site over its entire length was 1. Finally, we used FastTree (version 2.1.11, compiled for double precision; Price et al., 2010) to reconstruct unrooted maximum likelihood trees from these simulated alignments.

### Model validation

We ran our Bayesian sampling method on each of the 100 simulated trees for 2 *×* 10^4^ steps, discarding a burn-in of 2,000 steps and thinning the remaining chain down to 1,000 steps. We set the lognormal prior distribution on clock rates to *μ* = *−*5 and *σ* = 2, and the uniform prior distribution on root dates to a minimum of one simulation time unit before the true origin, and a maximum of the first HIV RNA sampling time. In addition, we set the proposal parameters to *δ* = 0.01 for the root location, *σ* = 0.33 for the time of infection, and *δ* = 0.01 for the clock rate. In preliminary runs, we found that these settings were sufficient for replicate chain samples to converge to the same posterior distribution. To sample integration dates for each DNA sequence, we further thinned the chain down to a total of 200 samples from the posterior distribution.

To compare our results against conventional root-to-tip regression, we censored the sampling times associated with tips that represented DNA sequences, and then rooted the tree using the *rtt* function in the R package *ape* (implementation by R. M. McCloskey; Paradis and Schliep, 2019). We extracted the root-to-tip distances from the resulting tree, and fit a simple linear regression of these distances against sampling times. Finally, we used the *inverse.predict* function from R package *chemCal* to extract predicted integration dates for the 200 samples from the posterior distribution.

To quantify the discordance between estimated 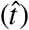 and actual (*t*) integration dates, we calculated the root mean square error, 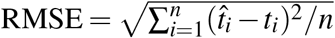, where *n* is the number of DNA sequences. We used a paired Wilcoxon rank-sum test to evaluate the significance of differences between the RMSEs obtained from *bayroot* and conventional RTT.

## 3. Results

To compare conventional root-to-tip regression (RTT) to our Bayesian approach (*bayroot*), we simulated the proliferation of HIV-1 among active and latent CD4+ T cells with an exact stochastic method. Our simulation workflow yielded a total of 100 trees reconstructed from HIV-1 RNA and DNA sequences. We assumed that HIV-1 RNA was sampled before the start of antiretroviral therapy (ART), and that HIV-1 proviral DNA was sampled from the latent reservoir in the post-ART period (Figure 2). 50 of the trees were simulated such that HIV-1 RNA was sampled at three time points starting at 3 time units post-infection, at intervals of three time units (scenario 1). For the remaining 50 trees, HIV-1 RNA sampling was delayed to 11 time units post-infection and taken at narrower intervals of two time units (scenario 2).

Figure 3 compares the estimates of HIV-1 DNA integration dates produced by RTT and (*bayroot*). Under scenario 1, both methods tended to produce similar estimates because the sampling conditions were favourable for fitting the molecular clock (Figure 3A). The median RMSE was 0.947 for RTT and 0.889 time units for *bayroot*. On a case-by-case basis, *bayroot* produced significantly more accurate estimates than RTT (paired Wilcoxon test, *P* = 3.55 *×* 10^*−*4^, Figure 3B). The overall difference between estimates was numerically small. For instance, the median difference in RMSE between RTT and *bayroot* was 0.059 (interquartile range, IQR = 0.004 *−* 0.201) time units. In some cases, however, integration dates were mapped by RTT to the time period after ART initiation, leading to higher RMSE values (Figure 3B). Since *bayroot* incorporates the prior information that HIV-1 integration should not occur during effective ART, its estimates are constrained to times preceding ART initiation. Furthermore, 89.8% of the actual integration dates fell within the 95% credible intervals generated by *bayroot*.

**Figure 3:**
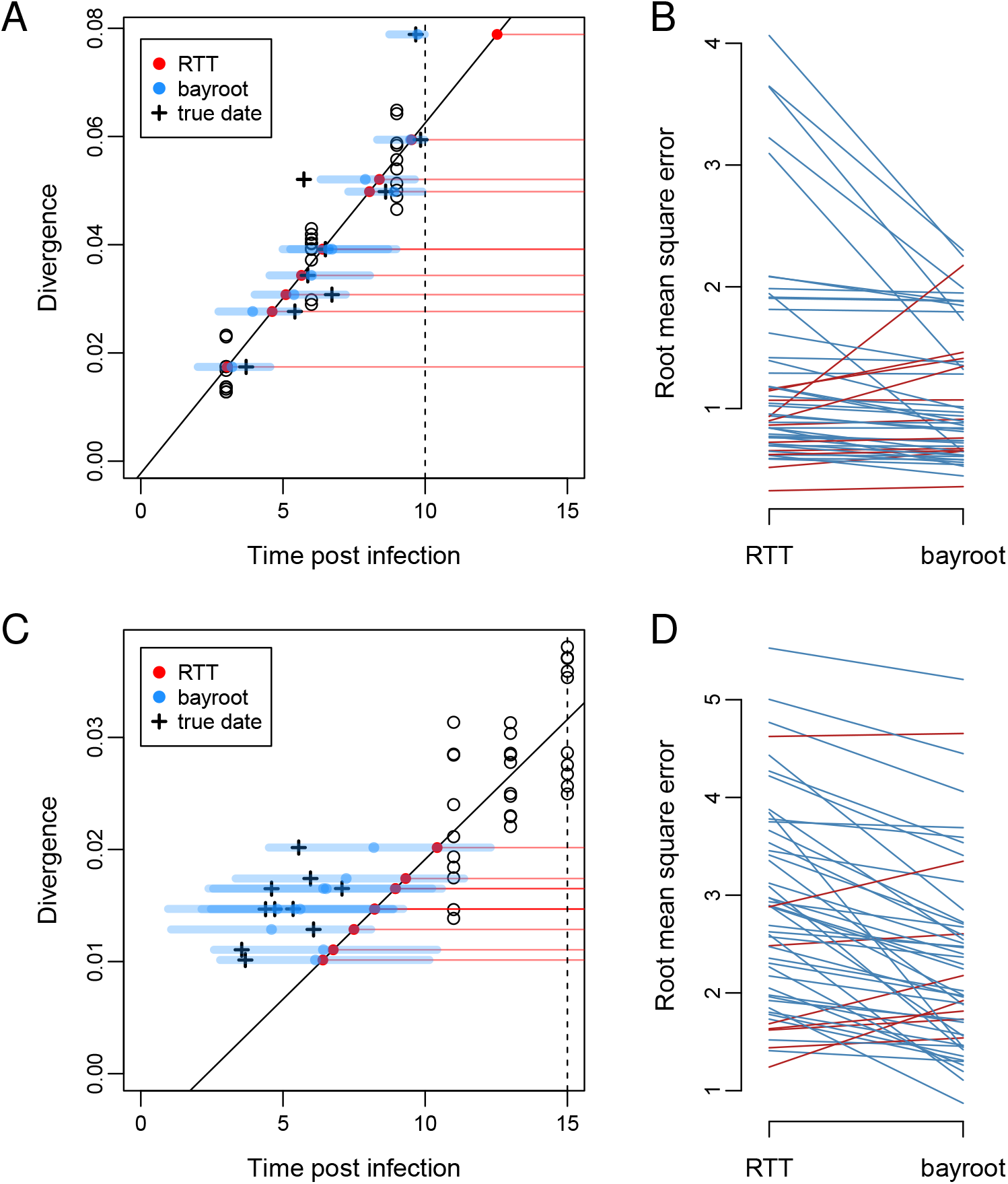
Comparison of results from *bayroot* and conventional root-to-tip (RTT) regression. (A) A scatterplot of root-to-tip distance (divergence) against sampling times post infection, for a representative example generated under scenario 1. A solid line represents the RTT regression fitted to the RNA sequence data (open circles), which we expect to intercept the horizontal axis at *t* = 0. A vertical dashed line marks the start of ART. Red points represent estimates of integration dates from the RTT model for DNA sequences sampled at time *t* = 20, as indicated by horizontal red lines. Blue points and line segments represent the median and 95% credible interval for integration date estimates from *bayroot*. Cross marks indicate the actual integration dates. (B) A slopegraph comparing the root mean square error (RMSE) of integration date estimates from RTT and *bayroot* for all 50 simulations generated under scenario 1. Line segments are coloured red if the RMSE for a given simulation was greater for *bayroot*, and blue otherwise. (C) and (D) A scatterplot and slopegraph for simulations generated under scenario 2. Slopegraphs was generated using R package *ggfree* (https://github.com/ArtPoon/ggfree).

For scenario 2, both methods became less accurate with median RMSEs of 2.79 and 2.10 time units for RTT and *bayroot*, respectively (Figure 3D). Because the sampling times of the RNA sequences used to calibrate the molecular clock were closer together and more distant from the actual time of infection in this scenario (Figure 3C), we are less certain about all three parameters of the regression, *i.e*., the location of the root in the tree, the time associated with the root (*x*-intercept), and the clock rate (slope). Under these conditions, *bayroot* benefits from having prior information about the time of infection. For our simulations where *t* = 0 is the actual time, we constrained the time of infection variable to the interval from *−*1 to 3 simulation time units. (In practice, one could use a uniform prior bounded by the last seronegative and first seropositive dates for that individual.) In other words, prior information about the time of infection ‘anchors’ the root-to-tip regression when there are insufficient data to accurately estimate the *x*-intercept (Figure 3C). As a result, *bayroot* was significantly more accurate than RTT (paired Wilcoxon test, *P* = 3.82 *×* 10^*−*7^, Figure 3D). The median difference in RMSE between RTT and *bayroot* was 0.405 (IQR 0.190 *−* 0.807) time units — about seven times greater than scenario 1. 89.4% of actual integration dates fell within the 95% credible intervals from *bayroot*. There was no significant association in this outcome between scenarios (Fisher’s exact test, odds ratio = 0.5, *P* = 0.34).

Running a chain sample for 2 *×* 10^4^ steps in *bayroot* required a median of 47.3 (IQR 45.0-48.8) seconds in R version 4.2.0 for Linux on a single core of an AMD Ryzen ThreadRipper 1950X processor.

## 4. Discussion

The reconstruction of HIV-1 integration dates is a challenging problem. Cells carrying replication-competent provirus in the latent reservoir comprise a small fraction of resting CD4+ T cells (approximately 0.01 to 10 per million cells; Crooks et al., 2015; Prodger et al., 2020). Sequences of plasma HIV-1 RNA or integrated DNA often cover only a portion of the virus genome (Laskey et al., 2016), making it difficult to resolve their evolutionary relationships. In addition, the development of phylogenetic and statistical methods for analyzing these sequence data (Ferreira et al., 2021) has lagged behind ongoing improvements in molecular techniques (Cho et al., 2022; Sun et al., 2022). Here we have described a Bayesian extension of a widely-used regression method for estimating HIV-1 integration dates from sequence variation in the latent reservoir (Brodin et al., 2016; Brooks et al., 2020; Jones et al., 2018). Our method provides a means of incorporating additional data about the infection — *e.g*., the estimated date of infection, time of ART initiation, and previous measures of the rate of HIV-1 evolution within hosts — as prior information. Furthermore, adopting a Bayesian approach enables us to quantify our uncertainty about parameter estimates by sampling from the posterior distribution. We expect this will be important for studies where there is limited access to longitudinal plasma samples for retrospective sequencing, for instance.

Of course, our method also retains some significant limitations of conventional approaches to root-to-tip regression. First, we are assuming that the unrooted phylogeny relating HIV-1 RNA and DNA sequences is known without error. It is possible to relax this assumption by adopting a hierarchical approach and replicating our regression analysis on a posterior sample of unrooted trees that may be generated by a Bayesian phylogenetic program such as MrBayes (Ronquist and Huelsenbeck, 2003) or BEAST (Drummond and Rambaut, 2007). This is less efficient than sampling from the joint posterior distribution of unrooted trees, substitution model, and the RTT regression parameters. Additionally, we are assuming that the divergence of each sequence is an independent outcome. This convenient approximation is clearly untrue because of identity by descent: sequences that share a more recent common ancestor will have a similar root-to-tip distance because they have inherited the same set of mutations. It is possible to overcome this limitation by adapting the covariance matrix of the regression model to the phylogenetic structure of the data (Neher, 2018).

Not all studies use root-to-tip regression to estimate HIV-1 integration dates. For example, one of the methods described by Abrahams et al. (2019) uses approximate maximum likelihood to reconstruct a host-specific phylogeny relating HIV-1 RNA and DNA sequences, and then locates the closest tip representing an RNA sequence for every tip representing a DNA sequence, which is assigned the sampling time of the RNA tip. Hence, the DNA sequences can only be associated with a finite number of integration dates. This approach benefits from extensive sampling of HIV-1 plasma RNA over the time period spanning the start of infection to ART initiation. If the ancestral HIV-1 RNA sequence most closely related to an HIV-1 provirus is not represented in the tree, then the latter would be mapped to another branch that may be associated with a sampling time that does not accurately estimate of the integration date. In contrast, RTT methods directly use the number of mutations carried by an individual DNA sequence to estimate its integration date. The other sequences are used to calibrate the linear model mapping this divergence to the timeline.

## 5. Data availability

*bayroot* is publicly available under the MIT license at https://github.com/PoonLab/bayroot. We have also provided the simulated data and R scripts used to perform the method validation and generate figures in this repository.

